# Bioinformatics Analysis of the Potentially Functional circRNA-miRNA-mRNA Network in Breast Cancer

**DOI:** 10.1101/2022.01.10.475557

**Authors:** Cihat Erdoğan, İlknur Suer, Murat Kaya, Zeyneb Kurt, Şükrü Öztürk, Nizamettin Aydın

**Author notes:** These authors contributed equally: Cihat Erdoğan, İlknur Suer, Murat Kaya.

## Abstract

**Objective:** Breast cancer (BC) is a heterogeneous type of cancer that occurs as a result of distinct molecular alterations in breast tissue. Although there are many new developments in treatment and targeted therapy for BC in recent years, this cancer type is still the most common one among women with high morbidity and mortality. Therefore, new research is still needed for biomarker detection.

**Methods:** GSE101124 and GSE182471 datasets were obtained from Gene Expression Omnibus (GEO) database to evaluate differentially expressed circular RNAs (circRNAs). The Cancer Genome Atlas (TCGA) and Molecular Taxonomy of Breast Cancer International Consortium (METABRIC) databases were used to identify the significantly dysregulated microRNAs (miRNAs) and genes considering the Prediction Analysis of Microarray (PAM50) classification. The circRNA-miRNA-gene relationship was investigated using the Cancer Specific CircRNA (v2.0) (CSCD), miRDB, miRWalk and miRTarBase databases. The circRNA–miRNA–mRNA regulatory network was constructed using Kyoto Encyclopedia of Genes and Genomes (KEGG) pathway and Gene Ontology (GO) annotation. The protein-protein interaction network was constructed by the STRING 2021 database and visualized by the Cytoscape tool (v3.9.0). Then, raw miRNA data and genes were filtered using some selection criteria according to a specific expression level in PAM50 subgroups. A bottleneck method was utilized to obtain highly interacted hub genes using cytoHubba Cytoscape plugin. The overall survival (OS) and disease-free survival (DFS) analysis were performed for these hub genes, which are detected within the miRNA and circRNA axis in our study.

**Results:** We identified three circRNAs, three miRNAs, and eighteen candidate target genes that may play an important role in BC. In addition, it has been determined that these molecules can be useful in the classification of BC, especially in determining the basal-like breast cancer (BLBC) subtype.

**Conclusions:** We conclude that hsa_circ_0000515/ miR-486-5p/ SDC1 axis may be an important biomarker candidate in distinguishing patients in the BLBC group, especially according to the PAM50 classification of BC.

## Introduction

Breast cancer (BC) is a heterogeneous type of cancer that occurs as a result of distinct molecular alterations in breast tissue (1). Circular RNAs (circRNAs) are evolutionarily conserved and stable RNA regulators that can behave as microRNA (miRNA) sponges, control the expression of target miRNAs, regulate alternative splicing mechanisms, and take an active role in the expression of the gene in which they are encoded (2). circRNAs have been shown to play crucial roles in the cell, and in recent years this RNA class has been one of the most important research focuses, particularly in the field of cancer (Fig. 1) (3) The precise biological classification of the BC subtype is critical for predicting the disease’s progression. Clinical management of BC is dependent on criteria such as tumor size, age, Estrogen (ER) and Progesterone (PR) expression, and the presence or absence of amplification and concurrent enhanced Human epidermal growth factor receptor 2 (HER2) expression. However, these indicators are currently insufficient for accurately categorizing individuals into sections with a high or low risk of relapse, as well as identifying subgroups resistant to therapy (4). Technological breakthroughs in recent decades have enabled molecular classification based on distinct global gene expression. mRNA expression patterns assessed using microarrays revealed that BC had distinct intrinsic fingerprints that may be utilized to classify tumors into intrinsic molecular subgroups (5). Comparisons of diverse gene signatures in BC have been investigated for a long time, and agreement in classification has been demonstrated to be a moderate-level in many instances (6-8). Despite considerable advances in this field, there is still a need for novel markers to refine categorization, particularly for some subtypes (4). The identification of new genes with variable expression across different types of BC, as well as the detection of miRNAs and circRNAs associated with these genes, may be critical for cancer categorization and potential treatment. Therefore in our study circRNAs that may be relevant to BC were detected using the GSE101124 and GSE182471 databases. MiRNAs with significantly altered expression in Prediction Analysis of Microarray (PAM50) subtypes were identified using The Cancer Genome Atlas (TCGA) and Molecular Taxonomy of Breast Cancer International Consortium (METABRIC) datasets. The TCGA dataset was also used to identify genes with dramatically changed expressions in BC. The circRNA-miRNA-Gene relationship was investigated using CSCD (9), miRDB (10), miRWalk (11) and miRTarBase (12) databases.

**Fig. 1.**
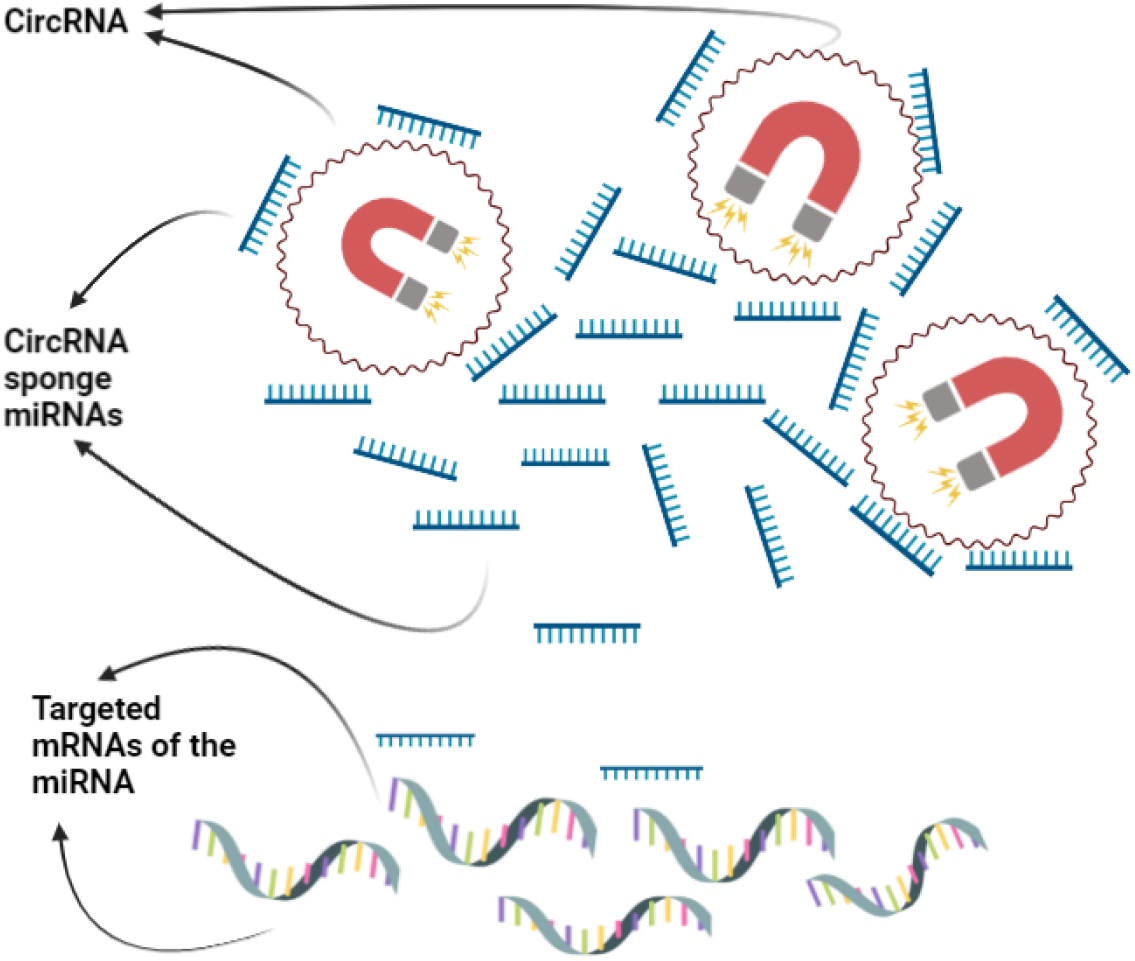
The diagram of circRNA-miRNA-gene relationship.

## Materials and Methods

### 1. Differentially expression analysis of BC datasets

circRNA, miRNA and gene expressions were analyzed using various databases and datasets. Block diagram of our pipeline is illustrated in Fig. 2.

a. **circRNA expression:** The circRNA expression profiles were collected from the Gene Expression Omnibus (GEO) database with an access code of GSE101124 (the dataset includes four BC cell samples, eight BC tissue samples and three mammary gland tissue samples) and GSE182471 (the dataset includes five BC samples and five non-tumor samples). The differentially expressed circRNAs (DECs) were identified by using the limma R package (v.3.46.0) with a p-value less than 0.05 and absolute log2-transformed FC (fold change) value of ≥1.
b. **miRNA and gene expression:** The miRNA and mRNA expression data as well as meta-data of the samples were downloaded from the TCGA database (13). The TCGA dataset contains 901 BC (162 basal-like, 73 HER2-positive, 455 Luminal A, 178 Luminal B, 33 normal-like) and 112 control miRNA and mRNA samples. Also, the METABRIC (University of Cambridge) dataset (14) was obtained from the European Genome-Phenome Archive (EGA-S00000000122) to validate our findings. The METABRIC dataset contains 1,301 BC (198 basal-like, 161 HER2-positive, 461 Luminal A, 370 Luminal B, 99 normal-like, and 12 unknown tumors) and 116 control miRNA samples. The details of the datasets are provided in supplementary file (S) Table S1. We used the DESeq2 R package (v.1.28.1) to identify differentially expressed mRNAs and miRNAs with a set of criteria, FDR < 0.01 and an absolute log2FC value of ≥1 (for both mRNAs and miRNAs), for each group of the PAM50 classification across the TCGA samples (Table S1) in a comparison to the healthy tissue samples. Since the raw data was not provided, but only the normalized data were available in the METABRIC database, the DEMs were determined with the same criteria given above by using the limma R package. Overlapping mRNA and miRNAs from differential expression analyses (i)-(v) were curated for further analysis. All FDR values were obtained by the Benjamini Hochberg method.

**Fig. 2.**
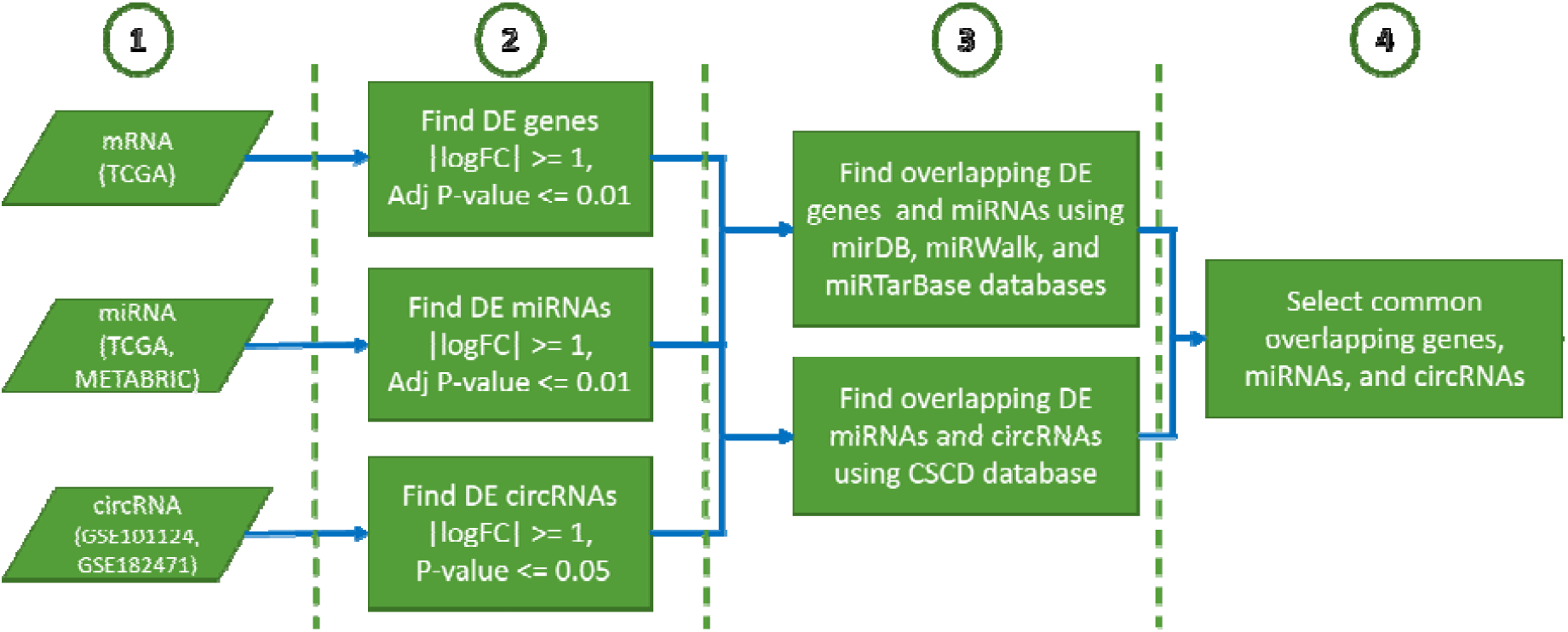
The steps of the BC PAM50 subtype analysis. The miRNA–mRNA interactions were predicted with mirDB (v6), miRWalk (v3), and miRTarBase (Release 8.0). DE: Differentially expressed, CSCD (v2.0): Cancer-specific circRNAs database, BC: Breast Cancer, FC: Fold change, log: logarithm base 2.

### 2. Predicting the associated biological features

In order to predict circRNAs and miRNAs interactions, the most significantly changed 16 DECs (4 down- and 12 up-regulated) (Fig. 3) were chosen via the CSCD v2.0 database. On the other hand, the miRDB, miRWalk and miRTarBase databases were used to find the interactions between the DEGs and DEMs across all of the PAM50 classes (Table 2). Hence, the knowledge base-driven target mRNAs of the up-regulated miRNAs in all PAM50 classes were searched among the down-regulated mRNAs, whereas the down-regulated miRNAs’ targets were searched among the up-regulated mRNAs. Similarly, knowledge base-driven target miRNAs of the up-regulated circRNAs were searched among the down-regulated miRNAs, whereas the targets of the down-regulated circRNAs were searched among the up-regulated miRNAs. After that, the interactions of circRNAs, miRNAs and mRNAs, which were found to have the most significant expression change, were investigated from the literature.

**Fig. 3.**
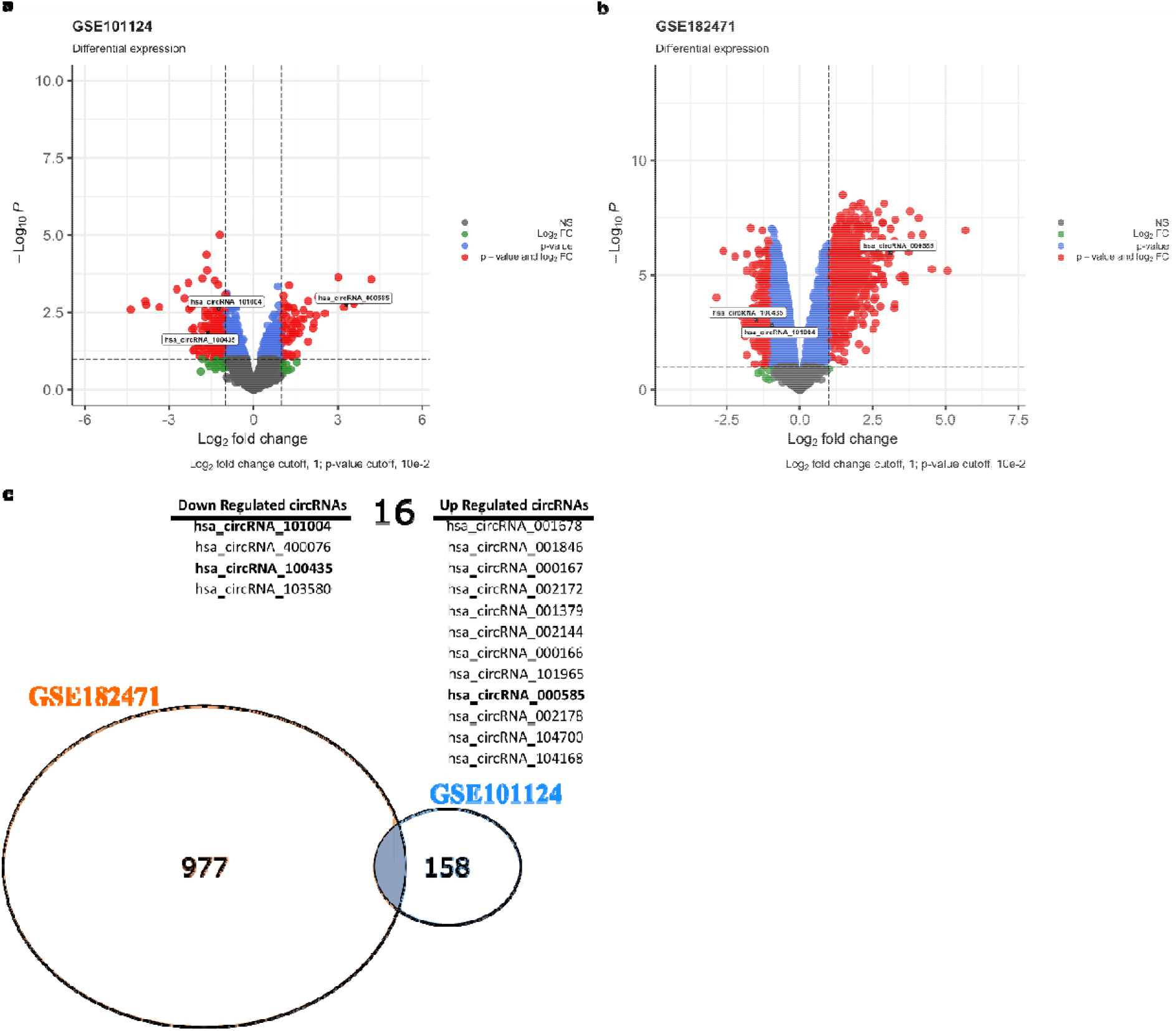
The volcano plot for DECs in BC based on the two microarray datasets from GEO and intersected up- and down regulated circRNAs. a: GSE101124, b: GSE182471, c: The intersected up- and down-regulated circRNAs between the GSE101124 and the GSE182471 datasets. DECs: differently expressed circRNAs. BC: Breast cancer.

### 3. Selection criteria of candidate miRNAs and genes

a. The selected miRNAs and genes should have a strong association with both BC and other cancers in the literature,
b. The selected genes should be associated with poor OS and/or DFS in BC (Fig. 9, 10, and 11),
c. A distinct altered expression level of these genes should be detected in PAM50 subgroups from normal-like to the BLBC (Fig. S-2a, 2b and 2c).

### 4. Survival analysis

The survival data of TCGA dataset was obtained from the Pan-Cancer Clinical publication (15). Survival curves were obtained according to the Kaplan-Meier method (surv_plot function from finalfit) (v1.0.3) R package (16), and differences between survival distributions were assessed by Log-rank test. The patients were divided into two risk groups as high and low according to their normalized median expression values. The normalized expression values obtained using *voom* function from *limma (v3*.*46*.*0) R package* (17). For analysis of relationships between the selected gene and BC, univariate models were fitted using Cox proportional hazard regression (*coxph* function from *survival R package* (18). Furthermore, we used the GSE25066 dataset from GEO to validate our survival analysis findings [cite]. The GSE25066 dataset contains 508 BC (189 BLBC, 37 HER2-positive, 160 LumA, 78 LumB, 44 normal-like) mRNA samples. The details of the clinical data for each datasets can be found in Table S1 separately.

### 5. The relation of circRNA, miRNA and mRNA

The circRNA–miRNA–mRNA regulatory network was constructed using *Cytoscape tool (v3*.*9*.*0)* (19) based on the interactions between circRNA, miRNA, and mRNA obtained from CSCD, mirDB, miRWalk, and miRTarBase databases.

### 6. Analysis of the protein–protein interaction (PPI) network

The PPI network was constructed by using the STRING 2021 (20) database and visualized by the *Cytoscape tool (v3*.*9*.*0)*.

### 7. Kyoto Encyclopedia of Genes and Genomes (KEGG) and Gene Ontology (GO) enrichment analyses

The gene set enrichment analyses were obtained by using *Enrichr* (23) web tool with the criterion of FDR value lower than 0.05 according to *GO* annotation and *KEGG* Pathway. *Enrichr* is a gene list enrichment analysis tool that is frequently used in the literature and allows querying on hundreds of gene sets such as *KEGG, GO, Reactome, DisGeNet*. The p-value, provided by *Enrichr*, as a result of the enrichment analysis is calculated by the Fisher’s exact test (hypergeometric test), which is a binomial proportionality test that assumes the binomial distribution and independence for the probability of any gene set. Also, the FDR value, provided by *Enrichr*, is calculated using the *Benjamini-Hochberg* method for correction for multiple hypothesis testing.

## Results

### Determination of DECs, DEMs and DEGs

a. **DECs:** We observed that 174 circRNAs (52 of them were up-regulated and 122 were down-regulated – given in Table S2) in GSE101124 (Fig. 3a) and 993 circRNA (665 of them were up-regulated and 328 were down-regulated – given in Table S3) in GSE182471 (Fig. 3b) in BC tumor samples were differentially expressed, when compared to the control samples. Furthermore, we obtained the overlapped down- and up-regulated circRNAs between the GSE101124 and GSE182471 datasets. The overlapped up- and down-regulated circRNAs are given in Fig. 3c and the details of these circRNAs are given in Table S4 and Table S5 for each dataset separately. The expression of the overlapped up- and down-regulated circRNAs in each dataset is shown in Fig. 4. The essential characteristics of the three circRNAs are displayed in Table 1. The basic structural patterns of the three circRNAs are given in Fig. 5. The unpaired two-samples wilcoxon test results according to tumor and control samples of the selected three DECs are given in violin plots in Fig. 6 for each circRNA dataset separately.
b. **DEMs:** Regarding the miRNAs, in TCGA dataset 133 miRNAs (66 of them were up-regulated and 67 were down-regulated) in BLBC samples, 114 miRNAs (65 up- and 49 down-regulated) in HER2-positive samples, 105 miRNAs (49 up- and 56 down-regulated) in LumA group, 133 miRNAs (69 up- and 64 down-regulated) in LumB group, and 78 miRNAs (41 up- and 37 down-regulated) in normal-like tumor group were differentially expressed, when compared to the control samples. The top 10 up- and down-regulated miRNAs are given in (Table S6) and 25 up-regulated (Table S7) and 17 down-regulated miRNAs (Table S8) were shared across all five PAM50 classes. In addition, the up- and down-regulated miRNAs are listed in Tables S9-S13 for each PAM50 subtype separately. Furthermore, in METABRIC dataset 69 miRNAs (34 of them were up-regulated and 35 were down-regulated) in BLBC samples, 66 miRNAs (31 up- and 35 down-regulated) in HER2-positive samples, 51 miRNAs (28 up- and 23 down-regulated) in LumA group, 70 miRNAs (31 up- and 39 down-regulated) in LumB group, and 31 miRNAs (17 up- and 14 down-regulated) in normal-like tumor group were differentially expressed, when compared to the control samples. The top 10 up- and down-regulated miRNAs are given in (Table S14) and 9 up-regulated (Table S15) and 10 down-regulated miRNAs (Table S16) were shared across all five PAM50 classes. In addition, the up- and down-regulated miRNAs are listed in Tables S17-21 for each PAM50 subtype separately. Finally, we obtained the overlapped DE miRNAs between the TCGA and METABRIC datasets from shared up and down miRNAs. The determined miRNAs are given in Fig. 7. The unpaired two-samples t-test results according to basal and control samples of the selected three (hsa-miR-141-5p, hsa-miR-183-5p, and hsa-miR-486-5p) DEMs are given as violin plot in Fig. 8 for each miRNA dataset separately.
c. **DEGs:** We observed that 5,143 genes (2,925 of them were up-regulated and 2,218 were down-regulated) in BLBC samples, 5,078 genes (2,442 up- and 2,636 down-regulated) in HER2-positive samples, 4,245 genes (2,066 up- and 2,179 down-regulated) in LumA group, 4,836 genes (2,325 up- and 2,511 down-regulated) in LumB group, and 2,850 genes (1,847 up- and 1,003 down-regulated) in normal-like tumor group were differentially expressed, when compared to the control samples. According to the criteria we created (criterion 3-a and 3-c), a distinct altered expression level of 18 genes (SDC1, PRAME, MELK, NEK2, EXO1, TPX2, BUB1, DLGAP5, CIDEC, ADH1B, TMEM132C, ACVR1C, LIPE, ABCA8, BTNL9, TNXB, GPAM and AOC3) were detected in PAM50 subgroups from the normal-like group to the BLBC group (supplementary file Fig. S-1, S-2).

**Fig. 4.**
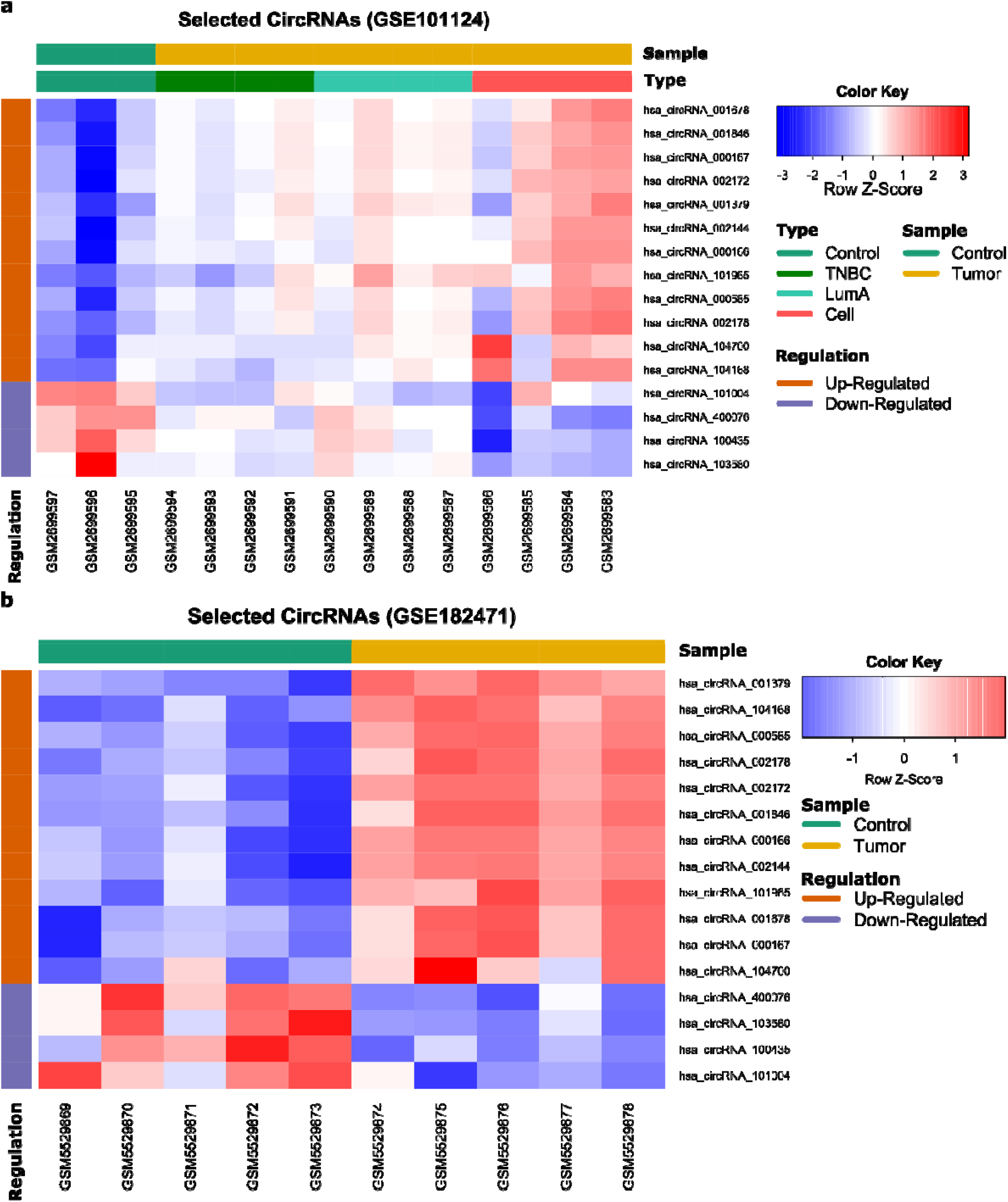
Heatmap for the overlapped up- and down-regulated DECs in individual microarray datasets, a: GSE101124, b: GSE182471. The heatmap was generated by R package ‘gplots’. DECs: differently expressed circRNAs.

**Table 1.**
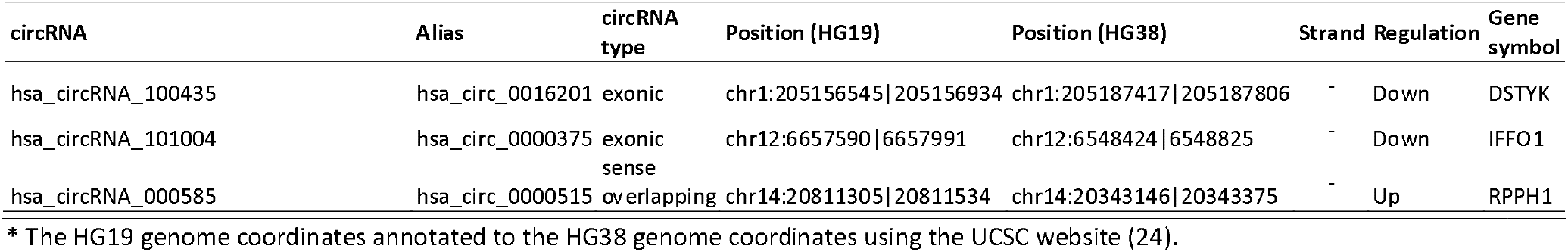
Essential characteristics of the three DECs.

**Table 2.**
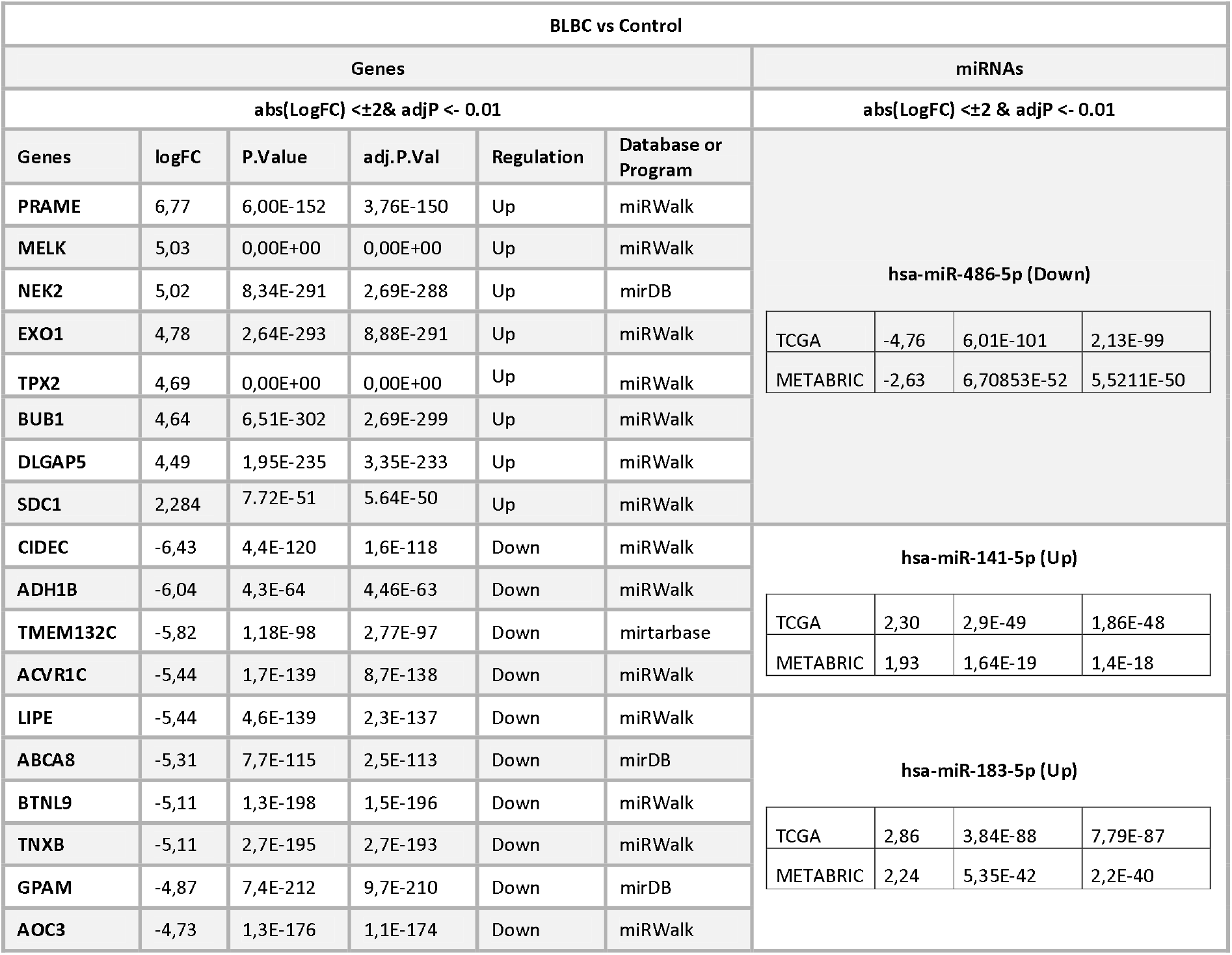
Expression levels of detected miRNA and possible target genes.

**Fig. 5.**
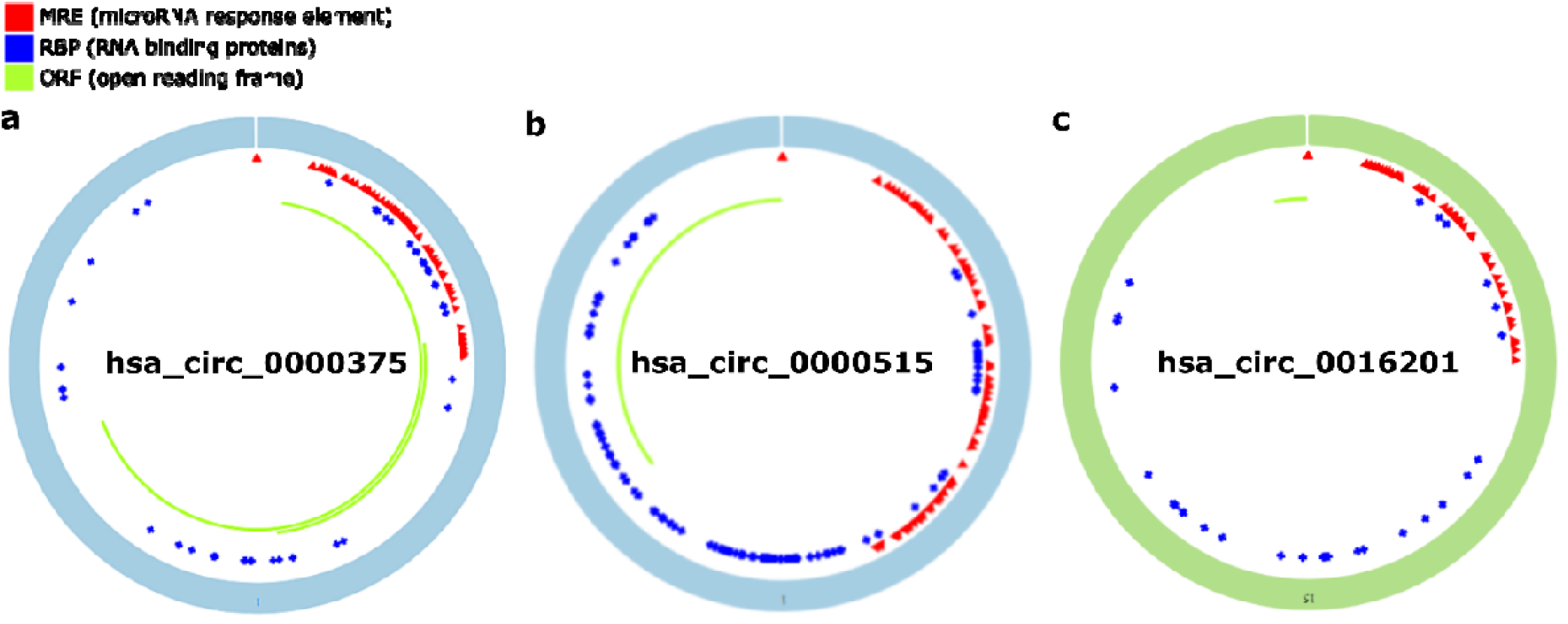
Structural patterns of the three DECs by the Cancer-Specific CircRNA (CSCD v2.0, http://geneyun.net/CSCD2/), **a:** has_circRNA_0000375, **b:** has_circRNA_0000515, **c:** has_circRNA_0016201. DECs: differently expressed circRNAs.

**Fig. 6.**
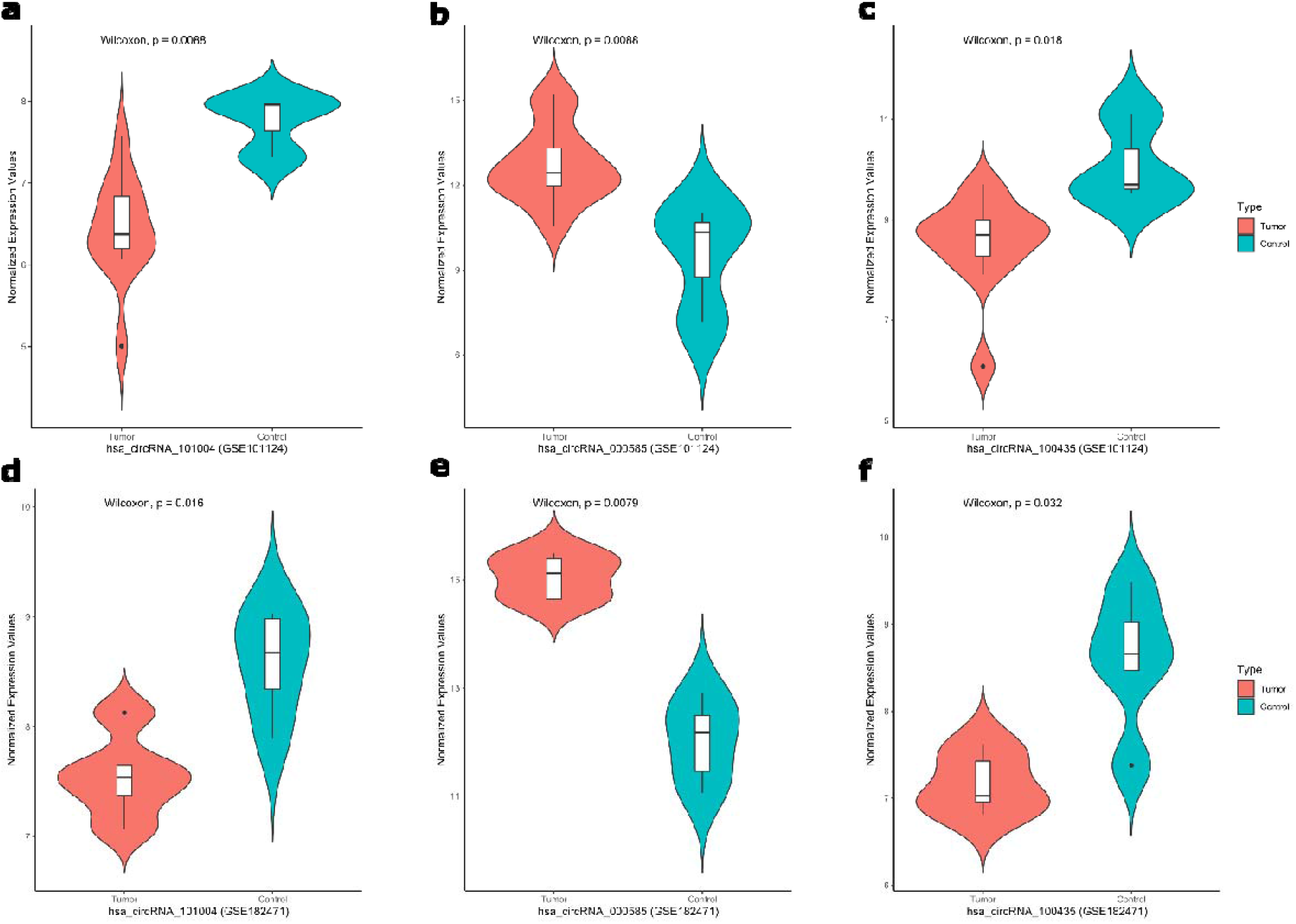
The combined violin and box plots for the normalized expression values of hsa_circRNA_101004, hsa_circRNA_000585, hsa_circRNA_100435 in GSE101124 and GSE182471 datasets by the unpaired two-samples wilcoxon test according to tumor and control samples. **a**: hsa_circRNA_101004 in GSE101124, **b:** hsa_circRNA_000585 in GSE101124, **c:** hsa_circRNA_100435 in GSE101124, **d**: hsa_circRNA_101004 in GSE182471, **e:** hsa_circRNA_000585 in GSE182471, **f:** hsa_circRNA_100435 in GSE182471.

**Fig. 7.**
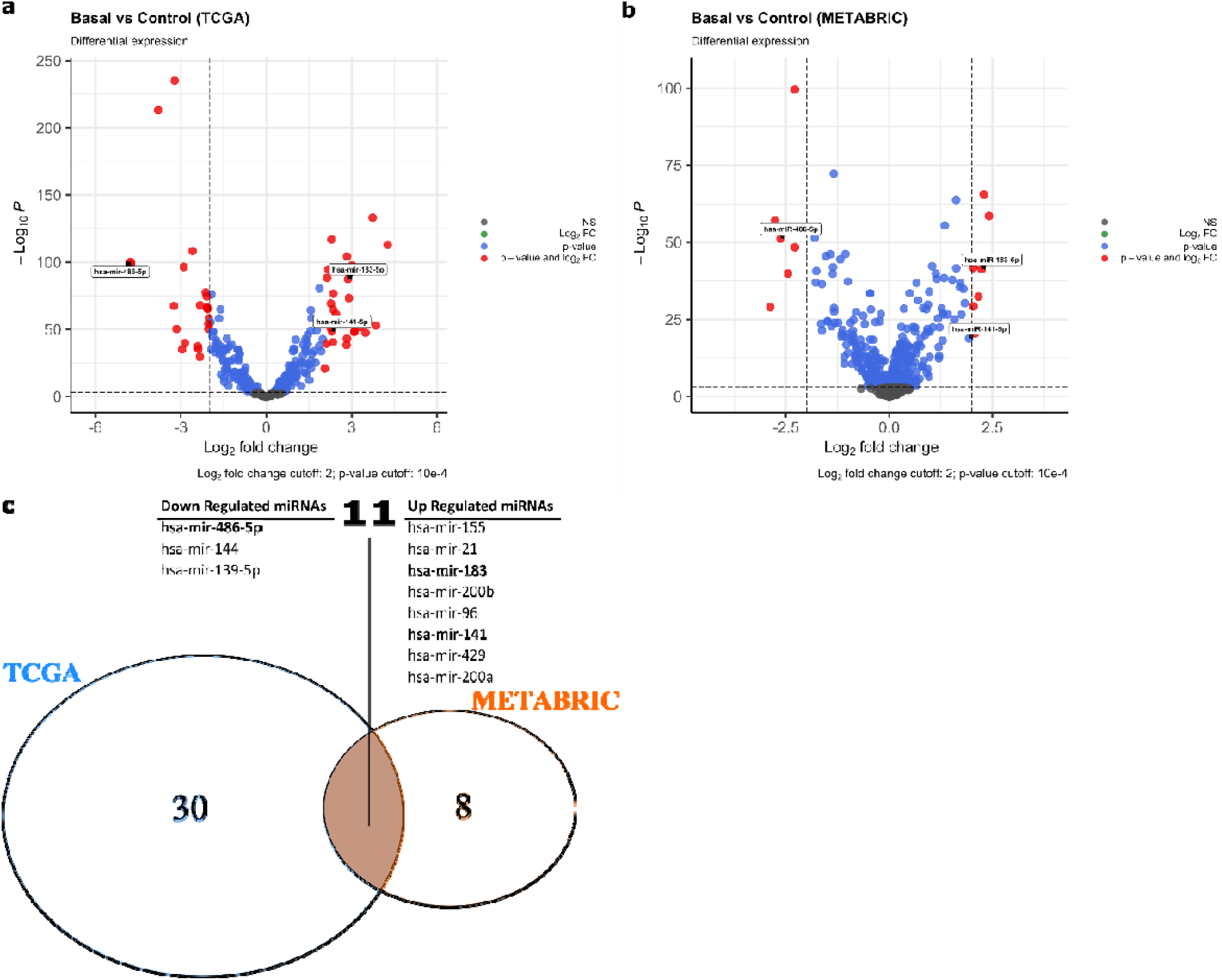
The volcano plot for DEMs in BC based on the two microarray datasets from TCGA and EGA, and intersected up- and down regulated miRNAs a: TCGA, b: METABRIC, c: The intersected up- and down-regulated miRNAs from shared miRNAs in the TCGA and the METABRIC datasets, DEMs: Differentially expressed miRNAs. EGA: European Genome-phenome Archive. BC: Breast cancer.

**Fig. 8.**
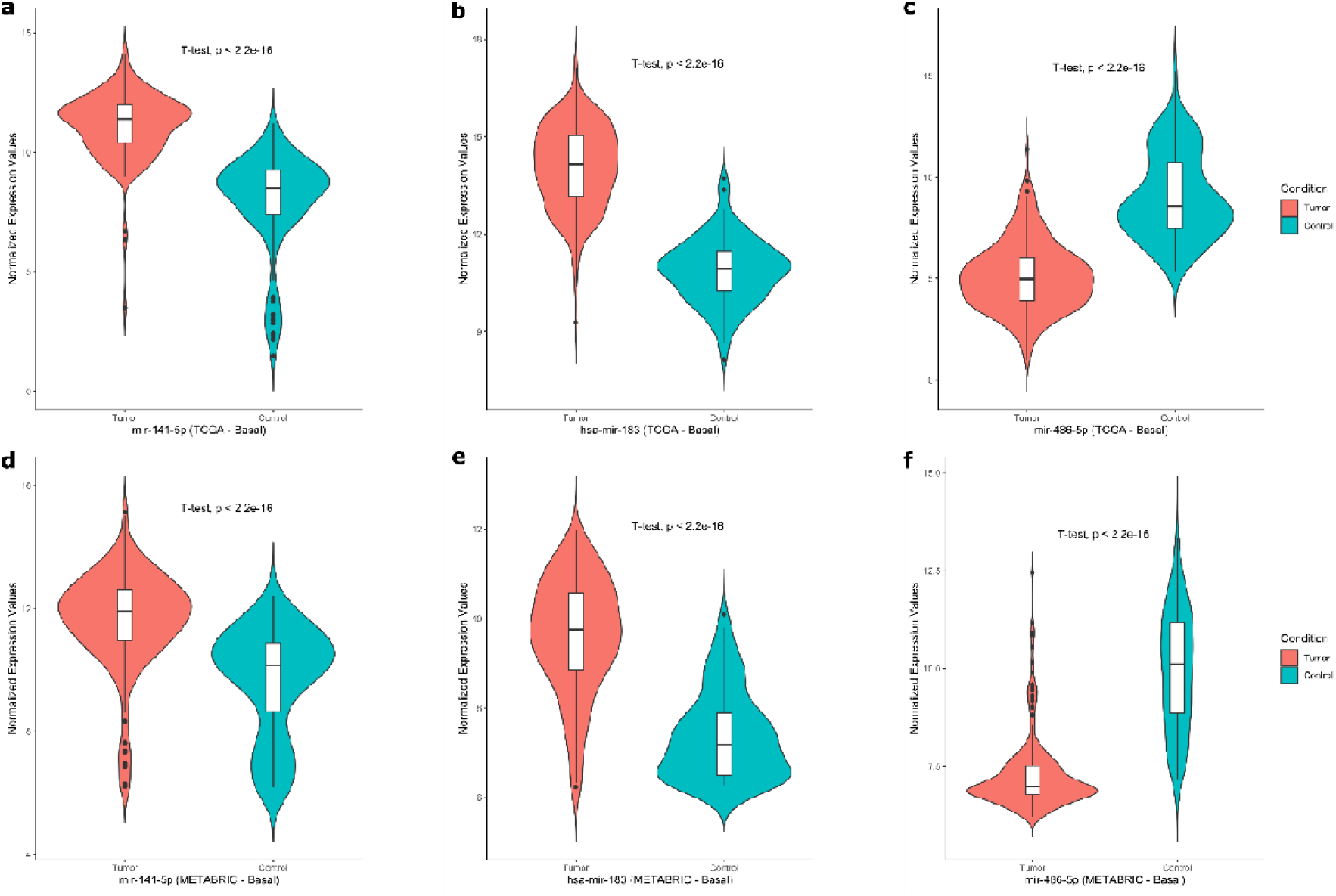
The combined violin and box plots for the normalized expression values of hsa-miR-141-5p, hsa-miR-183-5p, and hsa-miR-486-5p in TCGA and METABRIC datasets by the unpaired two-samples t-test according to basal and control samples. **a**: hsa-miR-141-5p in TCGA, b: hsa-miR-183-5p in TCGA, c: hsa-miR-486-5p in TCGA, d: hsa-miR-141-5p in METABRIC, e: hsa-miR-183-5p in METABRIC, f: hsa-miR-486-5p in METABRIC.

**Fig. 9.**
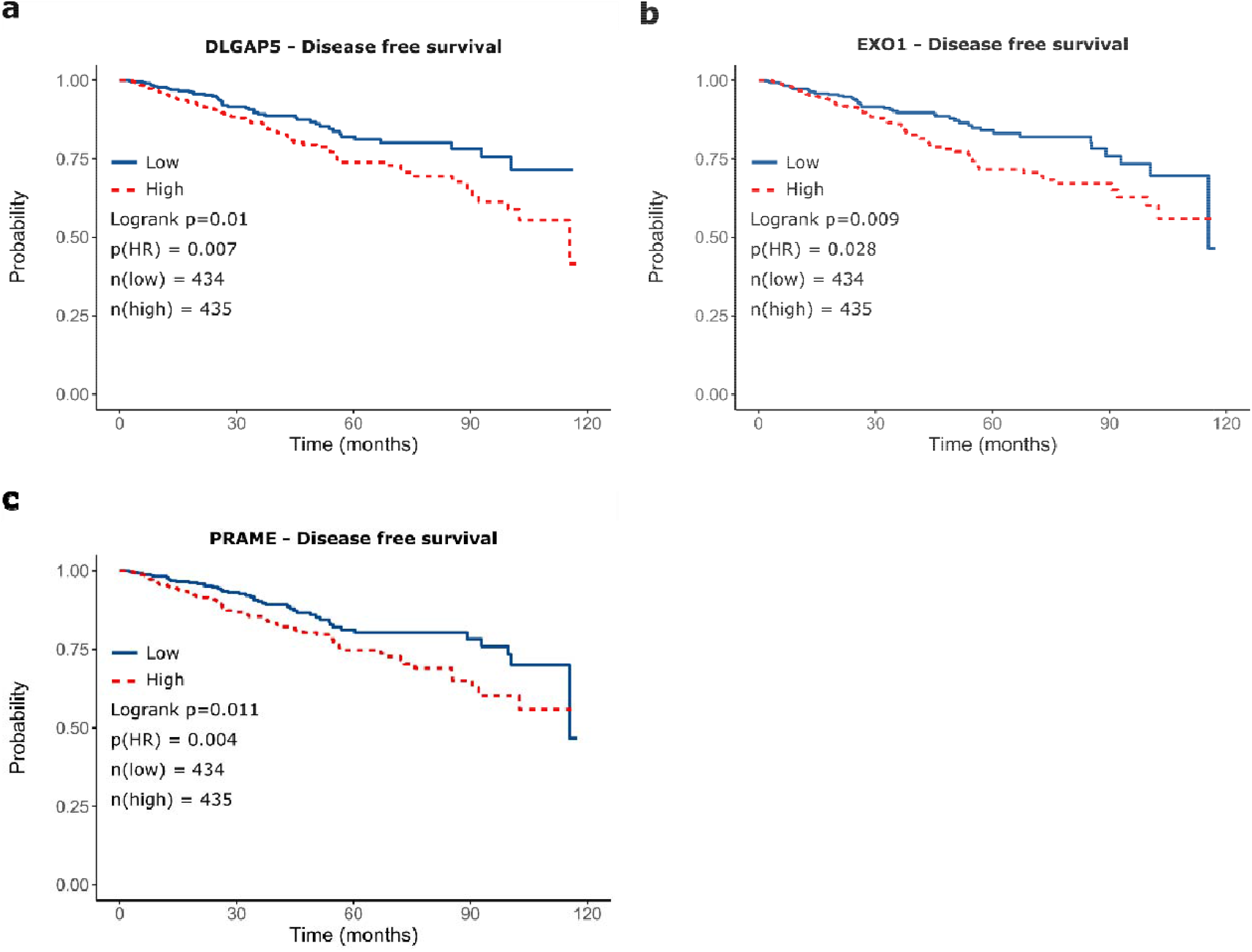
Survival analysis of selected genes from TCGA. Disease-free survival of genes a: DLGAP5, b: EXO1, c: PRAME.

**Fig. 10.**
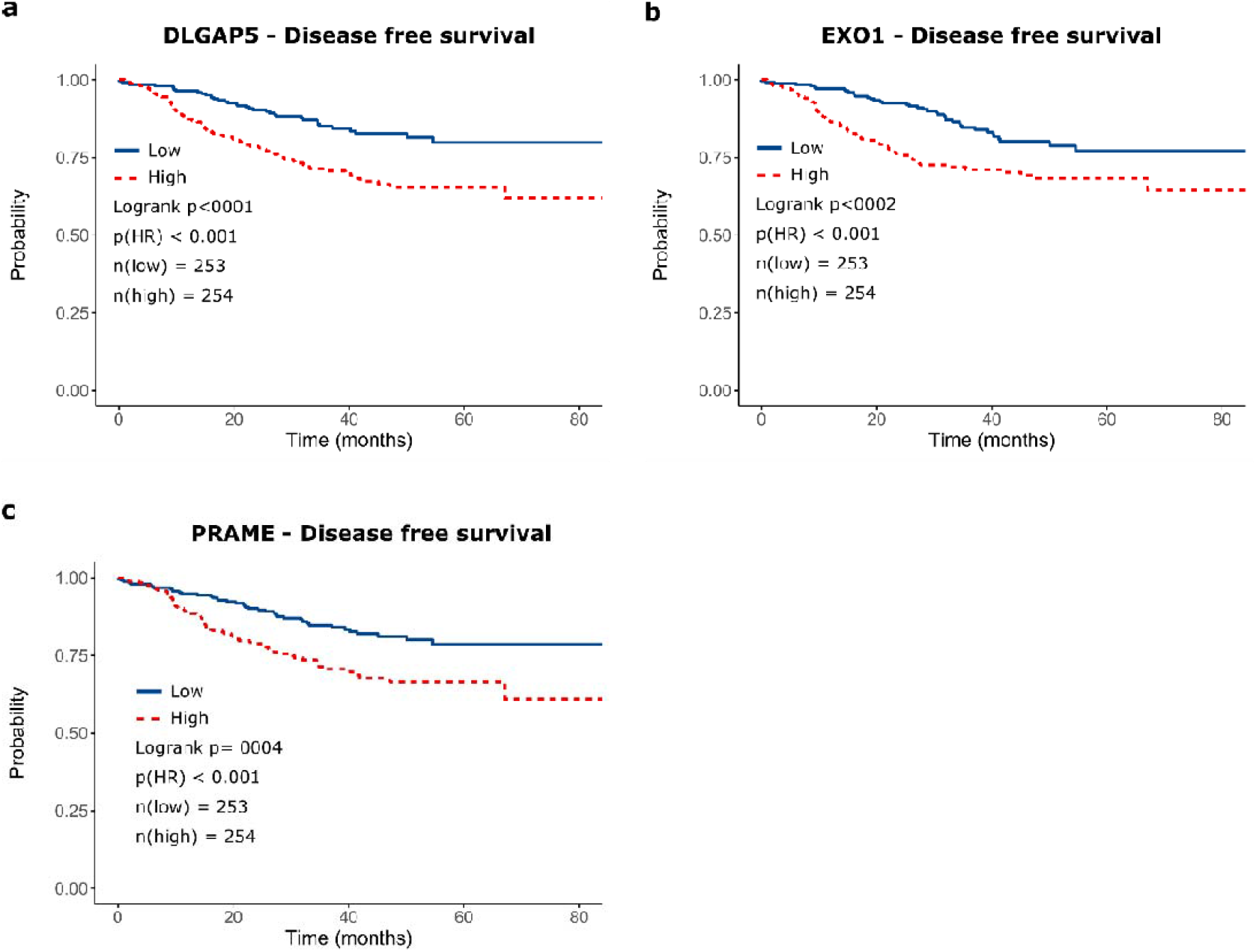
Survival analysis of selected genes from GSE25066 dataset. Disease-free survival of genes a: DLGAP5, b: EXO1, c: PRAME.

**Fig. 11.**
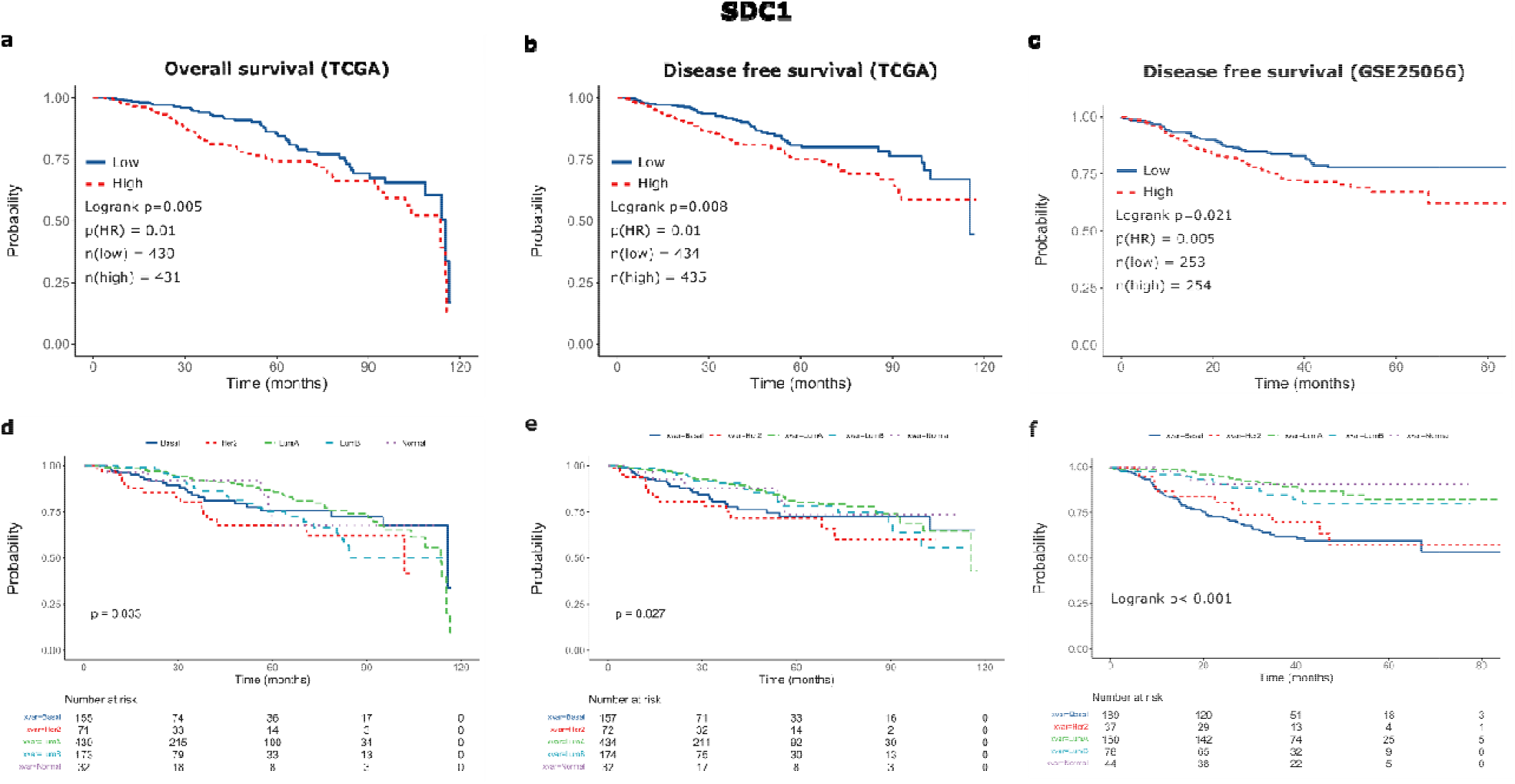
Survival analysis of SDC1 gene from TCGA and GSE25066 datasets. a: Overall survival of BC in TCGA, b: Disease-free survival of BC in TCGA, c: Disease-free survival of BC in GSE25066, d: Overall survival of PAM50 subgroups in TCGA, e: Disease-free survival of PAM50 subgroups in TCGA, f: Disease-free survival of PAM50 subgroups in GSE25066.

### Determining the relationship between the detected DECs, DEMs and DEGs

Knowledge-driven investigation between the shared mRNA and miRNAs, across all five PAM50 classes, revealed that 188 up-regulated genes are associated with the 3 down-regulated miRNAs; whereas 317 down-regulated genes are found to be associated to the 8 up-regulated miRNAs, based on the miRDB, miRWalk, and miRTarBase databases.

The top 10 up-regulated and down-regulated genes in each PAM50 class compared to the control samples according to our established criteria are listed in Table S22. Among the up-regulated genes, 746 were shared (Table S23) across all five PAM50 classes, whereas 650 down-regulated genes were shared (Table S24) across all five PAM50 classes. Additionally, the up- and down-regulated gene lists are provided in Tables S25-S29 for each PAM50 subtype individually.

### Survival Analysis

The increased expression of SDC1 gene had a poor overall survival (OS) and disease-free survival (DFS) and over-expressions of DLGAP5, PRAME, and EXO1 genes had a poor DFS (Fig. 9) in TCGA dataset. We also found that highly expressed SDC1 had a poor DFS, and over-expressions of DLGAP5, PRAME and EXO1 genes had a poor DFS (Fig. 10) in the GSE25066 dataset. The increased expression of SDC1 gene could provide a good estimation of poor OS and DFS in general BC and for all BLBC and HER2-positive subtypes (Fig. 11). The expression distributions of the SDC1, PRAME, EXO1 and DLGAP5 genes are shown in supplementary file Fig. S-3.

### Identification of the circRNA–miRNA interactions

The overlapped DECs were selected for further analysis. To depict whether the 11 circRNAs perform a significant role in BC, we collected their potential target miRNAs from the CSCD online databases. A total of three circRNA–miRNA interactions including three circRNAs (hsa_circRNA_000585, hsa_circRNA_101004, and hsa_circRNA_100435) and three miRNAs (miR-486-5p, miR-141-5p, and miR-183-5p) were identified. MIENTURNET (25) was exploited to explore the signaling pathways (*KEGG, Reactome, WikiPathways*, and *Disease Ontology*) in which the three miRNAs may be involved according to miRTarBase database. As shown in Fig. 12, all of the three miRNAs were related with some cancer-related pathways.

**Fig. 12.**
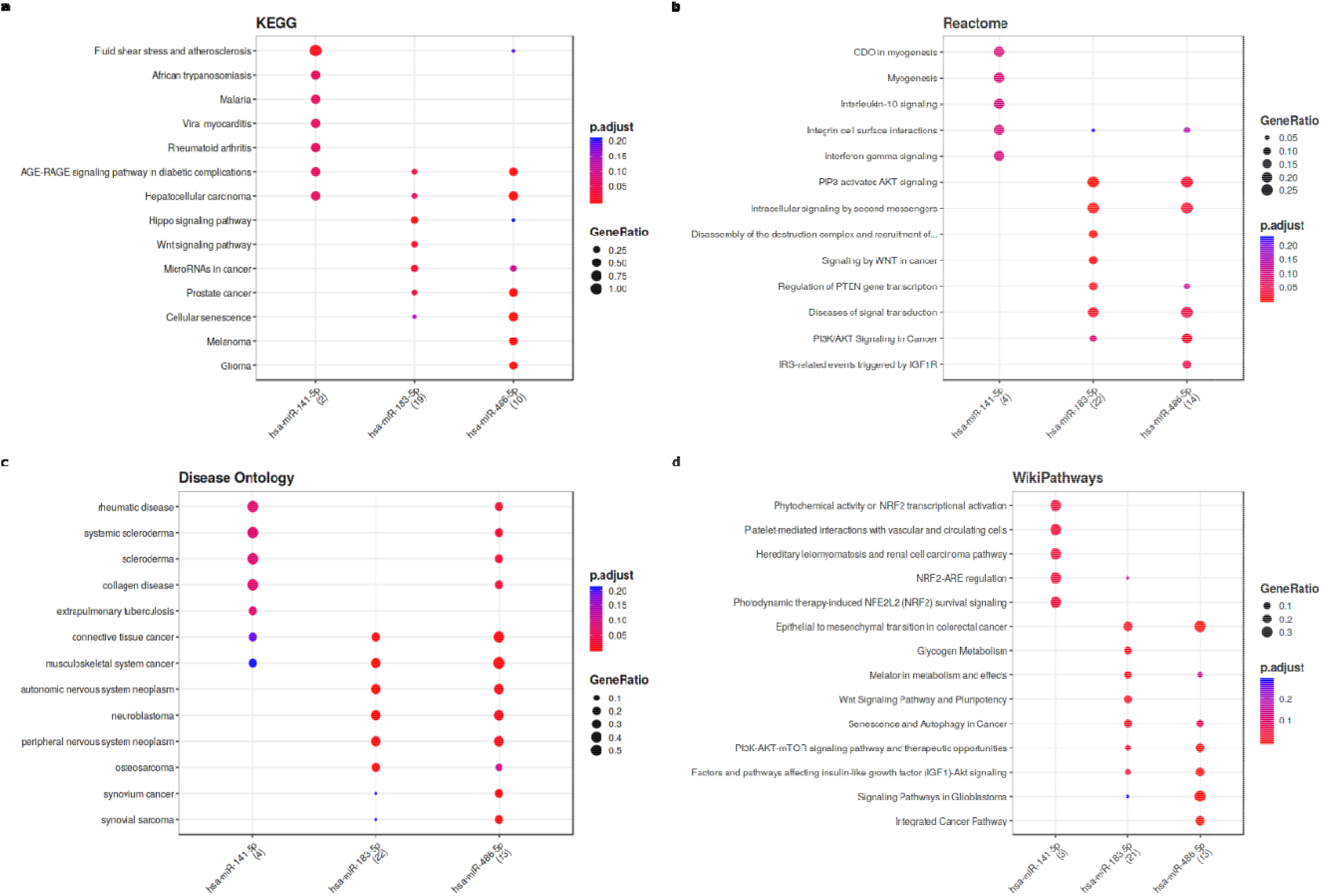
Enrichment analysis results for the significant signaling pathways that the three miRNAs related according to the MIENTURNET (http://userver.bio.uniroma1.it/apps/mienturnet/). **a:** KEGG pathways, **b:** Reactome pathways, **c:** Disease Ontology, **d:** WikiPathways.

### Identification of circRNA–miRNA–mRNA association

We investigated the miRNA and mRNA associations by using the miRDB (v6), miRWalk (v3), and miRTarBase (r8.0)] databases for the intersected miRNAs and shared DEGs with an absolute log2FC values greater and equal than 1. Then, we combined the circRNA–miRNA interactions and miRNA– mRNA interactions to identify the circRNA–miRNA–mRNA associations. Finally, we constructed a circRNA-miRNA-mRNA network, which provided a preliminary insight into the links between the three circRNAs (hsa_circRNA_000585, hsa_circRNA_101004, and hsa_circRNA_100435), the three miRNAs (miR-486-5p, miR-141-5p, and miR-183-5p) and the 339 mRNAs. The constructed network can be seen in Fig. 13.

**Fig. 13.**
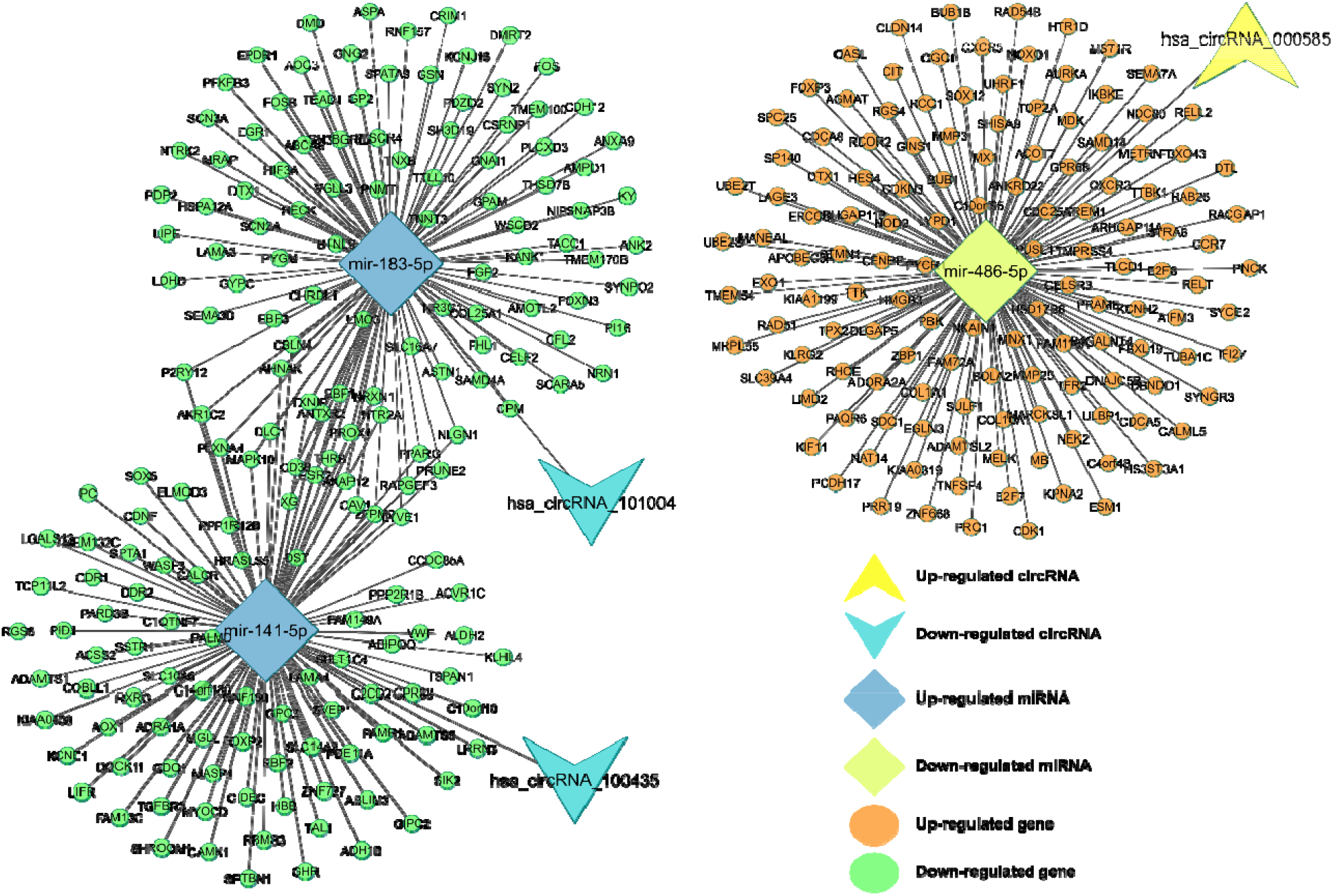
circRNA–miRNA–mRNA regulatory network. The network consisting of three circRNAs (hsa_circRNA_000585, hsa_circRNA_101004, and hsa_circRNA_100435), three miRNAs (miR-486-5p, miR-141-5p, and miR-183-5p) and 339 genes was generated by Cytoscape 3.9.0.

### Identification of the hub genes with bottleneck algorithm from the PPI network

Using the genes in Fig. 13, after removing the isolated nodes, we established a PPI network consisting of 138 nodes and 780 edges to view the interactions among the 339 mRNAs (Fig. 14a). Considering the importance of hubgene in a network, we employed a bottleneck algorithm to screen hub genes from the PPI network. The subnetwork with 14 nodes (10 hub genes and 4 extended genes) and 24 (14 between hub genes and 10 between extended genes) edges was identified (Fig. 14b), which unveiled the critical roles of the ten genes (AHNAK, CAV1, CDK1, EGR1, FGF2, FOS, KIF11, PPARG, SDC1, and TNXB) in BC. A circRNA-miRNA-hubgene network was then built to delineate the links among the DECs, DEMs and hub genes (Fig. 15). Thirteen circRNA–miRNA–mRNA regulatory modules, including hsa_circRNA_100435 / hsa-miR-141-5p / AHNAK regulatory axis, hsa_circRNA_100435 / hsa-miR-141-5p/PPARG regulatory axis, hsa_circRNA_100435 / hsa-miR-141-5p / CAV1 regulatory axis, hsa_circRNA_101004 / hsa-miR-183-5p/ AHNAK regulatory axis, hsa_circRNA_101004 / hsa-miR-183-5p / PPARG regulatory axis, hsa_circRNA_101004 / hsa-miR-183-5p / CAV1 regulatory axis, hsa_circRNA_101004 / hsa-miR-183-5p / FGF2 regulatory axis, hsa_circRNA_101004 / hsa-miR-183-5p / EGR1 regulatory axis, hsa_circRNA_101004 / hsa-miR-183-5p / TNXB regulatory axis, hsa_circRNA_101004/hsa-miR-183-5p/FOS regulatory axis, hsa_circRNA_000585 / hsa-miR-486-5p / CDK1 regulatory axis, hsa_circRNA_000585 / hsa-miR-486-5p / KIF11 regulatory axis, and hsa_circRNA_000585/ hsa-miR-486-5p / SDC1 regulatory axis, were found from the network.

**Fig. 14.**
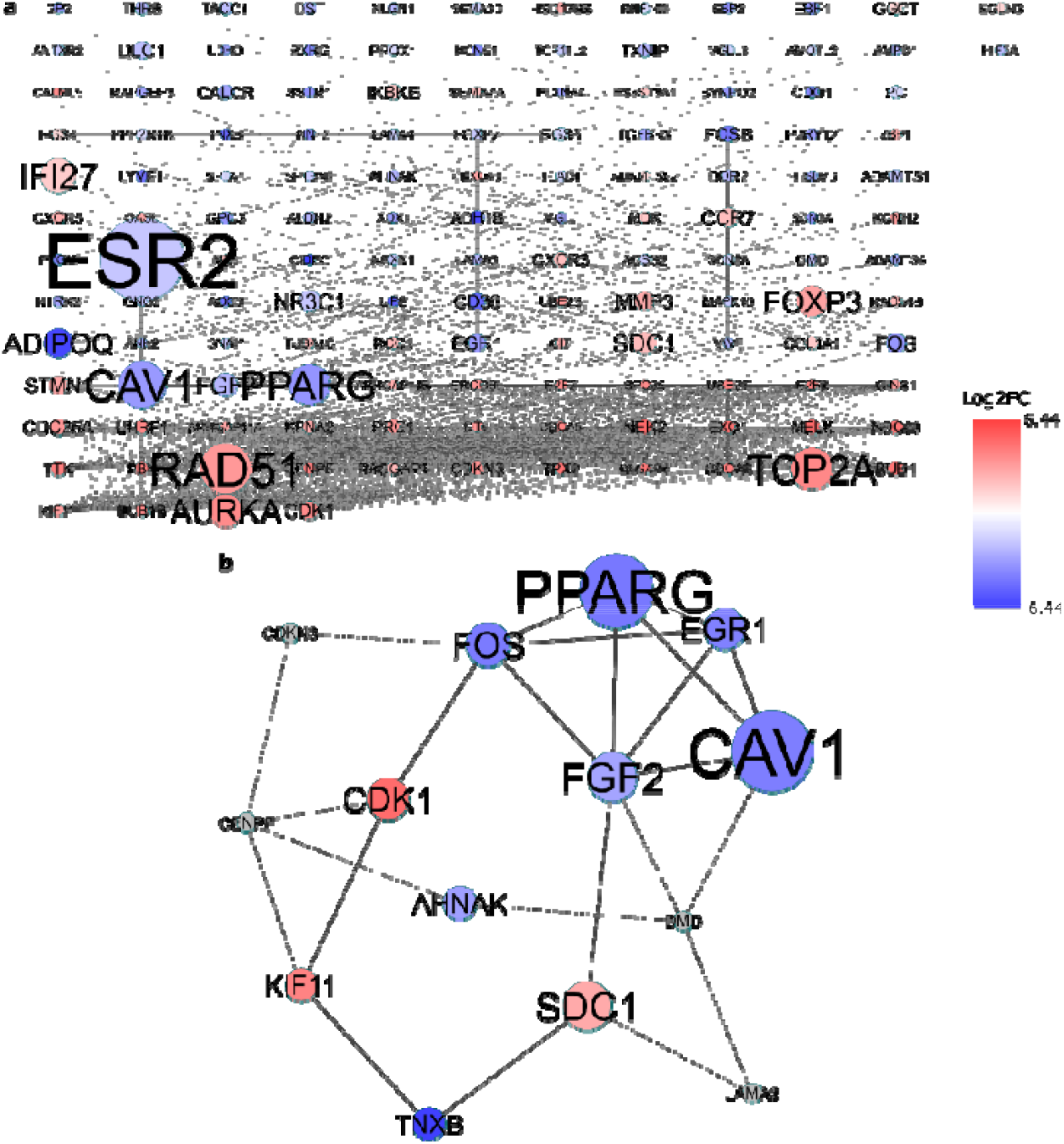
Identification of hub genes from the PPI network with the bottleneck algorithm using the cytoHubba Cytoscape plugin. The node color changes gradually from blue to red in ascending order according to the log2 (fold change) of genes. a: A PPI network of the 339 target genes that exert momentous roles in BC. This network consists of 138 nodes and 780 edges. The node size changes gradually from small to large in ascending order according to the number of the PMIDs from DisGeNET per gene. b: A PPI network of the 10 hub genes (colored blue and red) and 4 extended genes (colored gray) that extracted from a. This network consists of 14 (10 hub genes and 4 extended genes) nodes and 24 (14 between hub genes and 10 between extended genes) edges. PPI protein–protein interaction, BC: Breast Cancer.

**Fig. 15.**
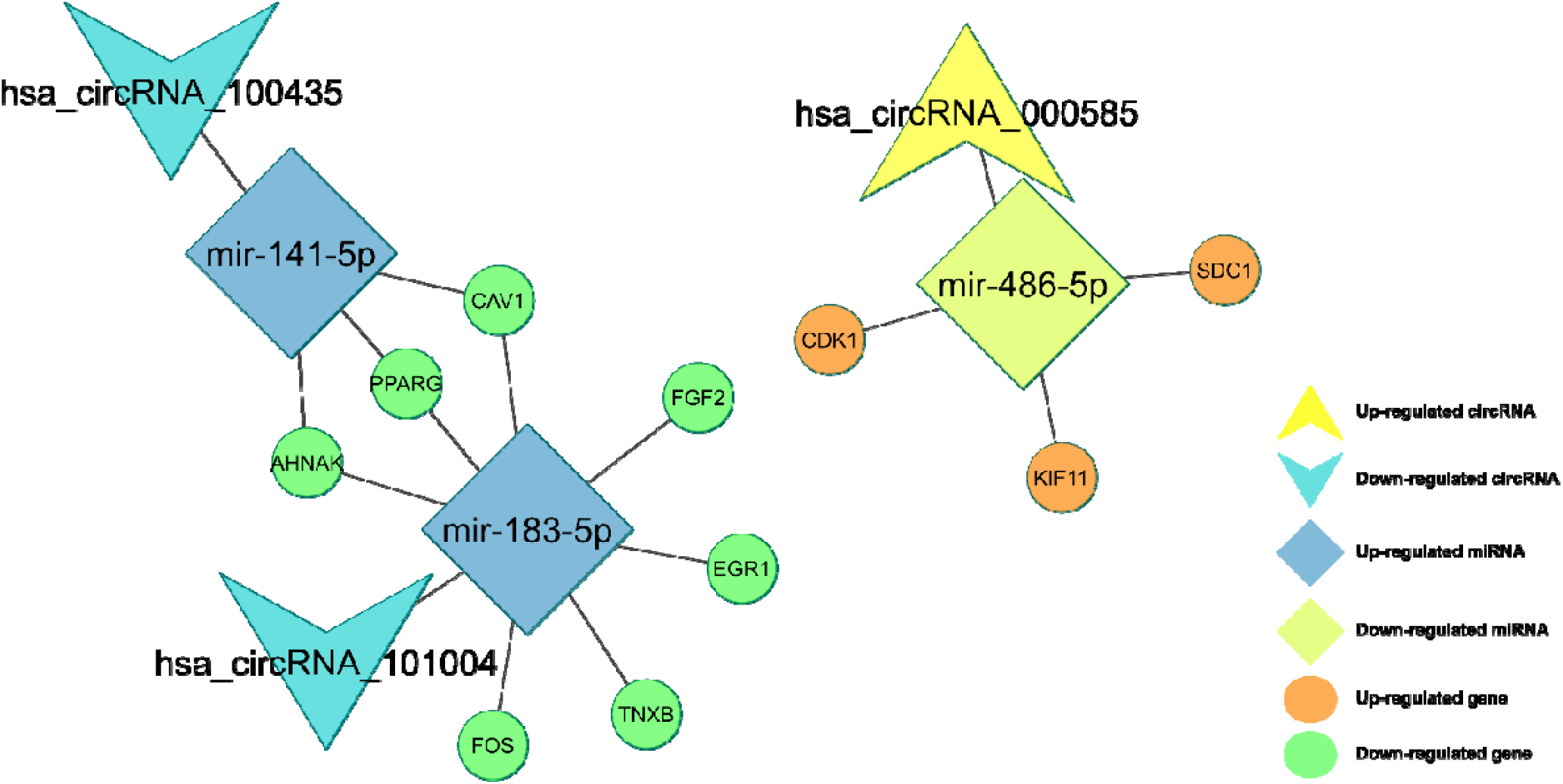
CircRNA–miRNA–hubgene network. The network consisting of three circRNAs (hsa_circRNA_000585, hsa_circRNA_101004, and hsa_circRNA_100435), three miRNAs (miR-486-5p, miR-141-5p, and miR-183-5p) and 10 hub genes (AHNAK, CAV1, CDK1, EGR1, FGF2, FOS, KIF11, PPARG, SDC1, and TNXB) was generated by Cytoscape 3.9.0.

### GO annotation and KEGG pathway analyses of the ten hub genes

GO analysis was carried out to illustrate the functional annotations of the ten hub genes. The top five highly enriched GO terms of biological process (BP), cellular component (CC) and molecular function (MF) are shown in Fig. 16. The most enriched GO terms in BP was “positive regulation of pri-miRNA transcription by RNA polymerase II (GO:1902895)” (FDR = 7.56E-07), that in CC was “sarcolemma (GO:0042383)” (FDR= 8.51E-03), and that in MF was “transcription regulatory region nucleic acid binding (GO:0001067)” (FDR= 7.74E-03). KEGG pathway analysis was conducted to ascertain the signaling cascade that the ten genes participate in. With an FDR < 0.05, 17 significantly enriched pathways were obtained (Fig. 17). Among the 17 pathways, “Proteoglycans in cancer pathway” and “Breast cancer pathway” are linked with the progression of BC (26, 27). Additionally, some other pathways such as “Pathways in cancer”, “Chemical carcinogenesis”, and “Non-alcoholic fatty liver disease” were also tumor-related pathways.

**Fig. 16.**
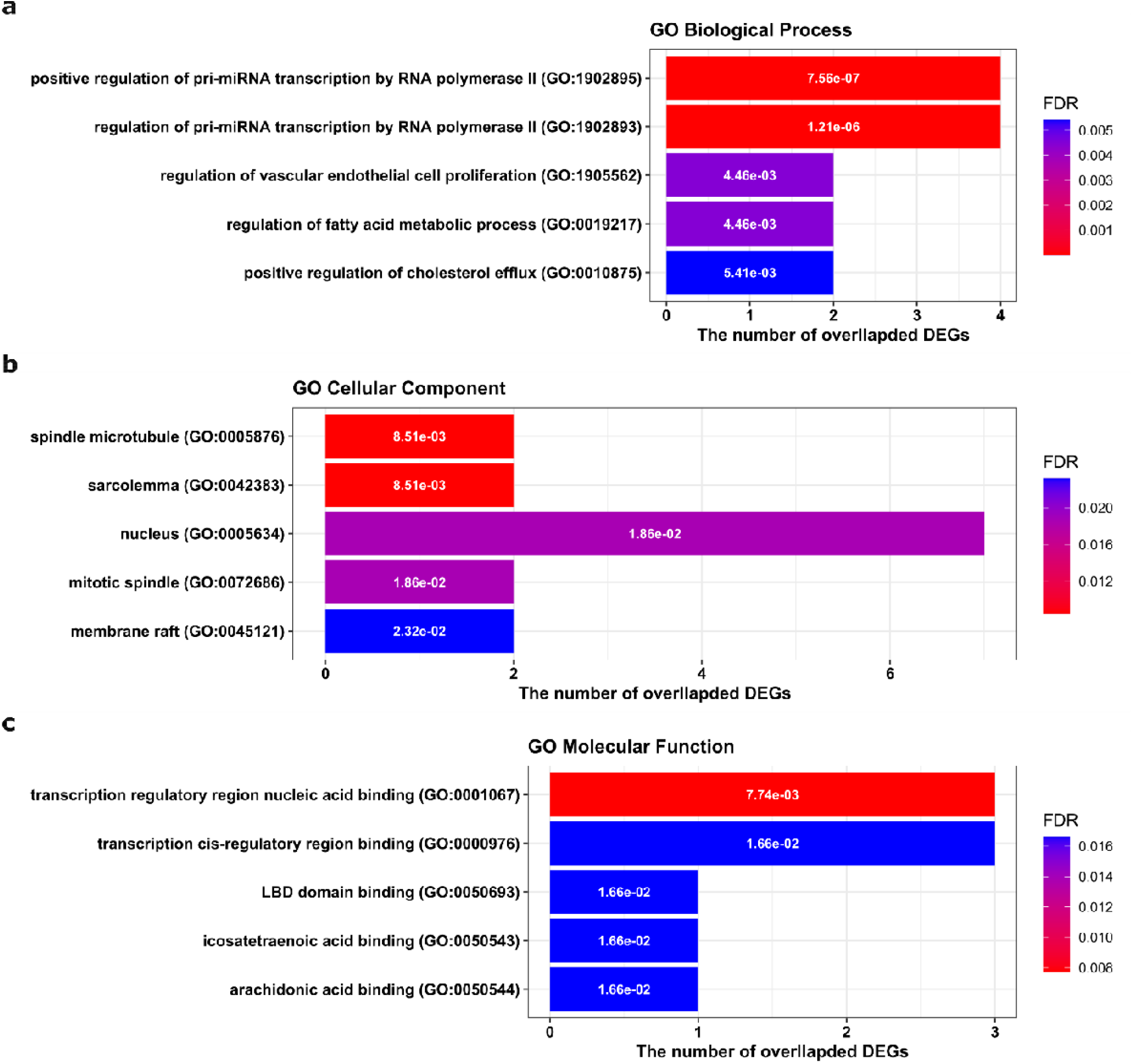
Top five Gene Ontology (GO) enrichment annotations of the ten hub genes: **a:** biological process, **b:** cellular component, **c:** molecular function. GO analysis was conducted by the ‘Enrichr’ web tool (https://maayanlab.cloud/Enrichr/) and visualized by R package ‘ggplot2’. DEGs: Differentially expressed genes. FDR: False discovery rate, is an adjusted p-value calculated using the Benjamini-Hochberg method for correction for multiple hypotheses testing.

**Fig. 17.**
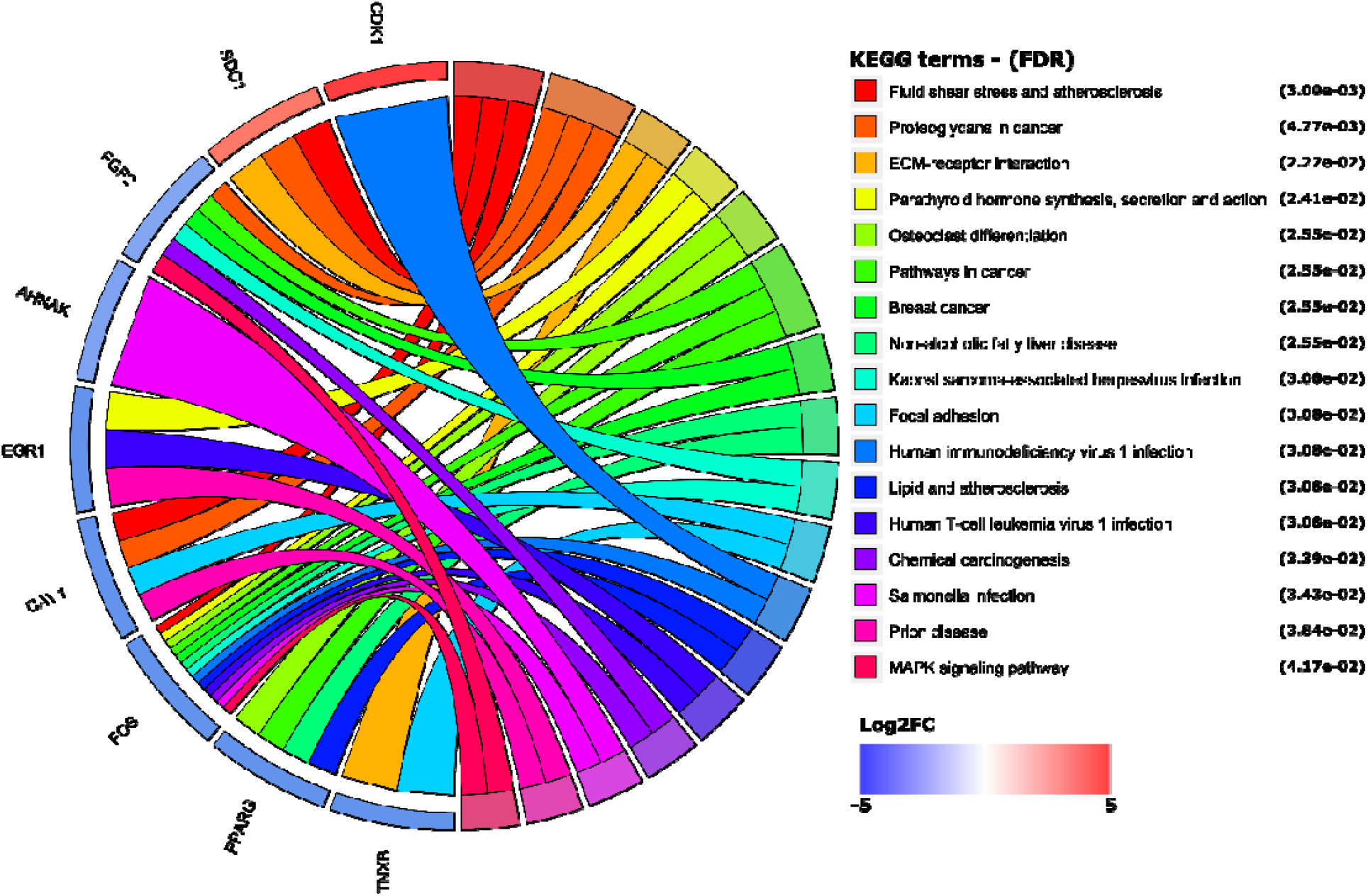
The significantly enriched Kyoto Encyclopedia of genes and genomes (KEGG) pathways with a FDR < 0.05. KEGG analysis was conducted using the ‘Enrichr’ web tool (https://maayanlab.cloud/Enrichr/) and visualized by R package ‘ggplot2’. Cohort plot shows that the ten hub genes are correlated via ribbons with their assigned KEGG terms. FDR: False discovery rate, is an adjusted p-value calculated using the Benjamini-Hochberg method for correction for multiple hypotheses testing.

### Filtered circRNAs, miRNAs and genes

The circRNAs with Log2FC value ≥1 (hsa_circ_0000515, hsa_circ_0016201, hsa_circ_0000375) were selected in both datasets (GSE101124 and GSE182471). 3 miRNAs (hsa-miR-486-5p, hsa-miR-141-5p, hsa-mir-183-5p) with Log2FC value ≥2 in both datasets (TCGA and METABRIC), which was reported to be strongly associated with BC and could be sponged by the circRNAs (detected via CSCD) were selected. Then, the 18 genes with a Log2FC value of ≥2 for the comparison of BLBC versus control samples (in PAM50 groups) in TCGA dataset which was reported to be strongly associated with BC, and could be targeted by the miRNAs, were determined. The possible circRNA-miRNA-mRNA interaction, which was detected to play a role in cellular processes of BC, is shown in Fig. 18.

**Fig. 18.**
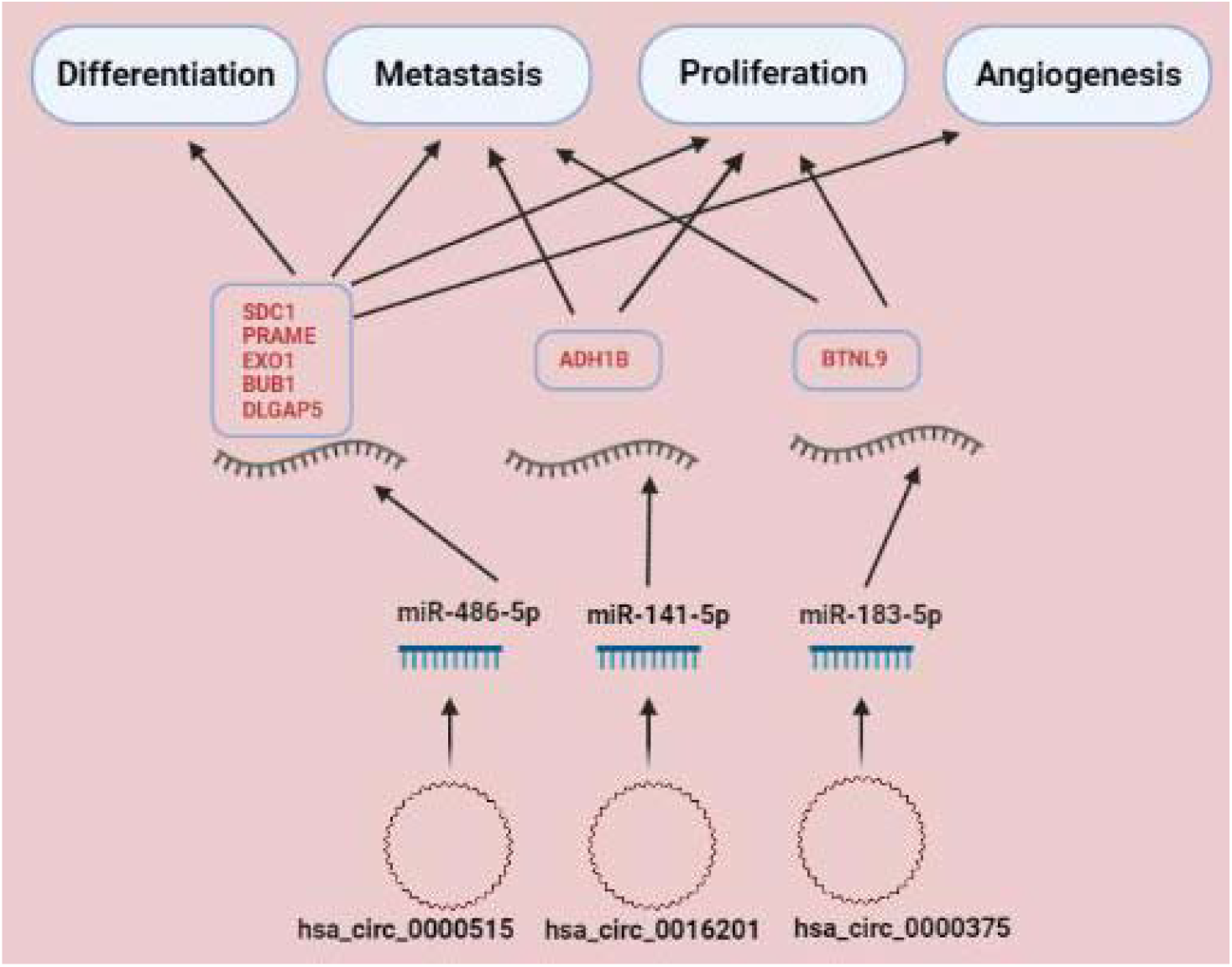
The summary of the possible role of circRNA/ miRNA/ gene axis in BC pathogenesis according to our study.

## Discussion

BC is the most frequently identified tumor among women around the world and more than 90% of BC deaths are related to metastasis. Existing treatment approaches for metastatic BC have been inadequate, compounded by a lack of early prognosis/ predictive criteria for estimating which body parts are most susceptible to metastasis. Although there are many new developments in the fields such as chemotherapy, endocrine treatment and targeted therapy for BC in recent years, this cancer type is still the most common cancer in women with high morbidity and mortality (28). Subtypes in BC are heterogeneous and treatment practices are determined according to these subtypes. From better to the worst, the aggressiveness of the BC subgroups are generally in the following order: Normal breast-like, LumA, LumB, HER2-positive, BLBC (29, 30). It is clearly known that the OS of cases with HER2-positive and BLBC groups are worst in PAM50 subtypes (31). BLBC is a highly aggressive molecular subgroup. The cells are “basal-like,” that implies they match the basal cells which line the breast ducts. It is strongly associated with triple-negative breast cancer (TNBC) appearance described by the absence of expression of ER, PR, and HER2-positive. BLBC, which is more associated with distant metastasis, has an extremely poor prognosis compared to other intrinsic BC groups, and the success in its treatment is currently limited (32, 33). This knowledge has substantially advanced our understanding of BC’s heterogeneity and the several biological processes that the disease employs.

In 2009, Parker et al. defined a minimum gene set, PAM50, for categorizing these intrinsic subgroups (34). Because the biology of all these intrinsic subgroups indicates changes in incidence, responsiveness to therapy and survival, unique genes for each subtype may be evaluated as markers to direct potential treatments. In this respect, it is crucial to elucidate novel gene-miRNA-circRNA relationships in the determination of these subgroups (35-37). In our study, the expression states of miRNAs and genes in datasets were classified according to the five molecular subtypes classification.

circRNAs, which are a new class of endogenous evolutionarily conserved RNAs, have a stable structure and they are stated to serve as vital regulators in the various cellular activities. According to studies conducted so far, it has been understood that the main reason for circRNAs to act as critical regulators in cells is their relationship with target miRNAs (38). In recent years, it has been suggested that miRNAs act as a bridge in the realization of the role of cirRNAs in the regulation of cellular processes (39). CircRNAs change gene expression by acting as miRNA sponges with their binding sites (3, 40). As increased expression rates of circRNAs in the cell may contribute to decreased expression levels of target miRNAs and the increased expression levels of target genes. circRNA-miRNA-mRNA interactions, which are the focus of this work, are very new to the scientific world but experiments have shown that these relationships could be beneficial for detection of novel biomarkers for cancer (41, 42). The studies on circRNAs about the determination of sub-types of BC are limited. The study by Nair et al. in 2016 is one of the first studies showing circRNAs may be useful in identifying subtypes of BC (43). In the study of Darbeheshti et al. in 40 TNBC, 20 Lum A, 18 Lum B and 17 HER2-positive tumor samples, it was determined that hsa_circ_0044234 has a distinct molecular signature as a potential GATA3 regulator in TNBC (44). In another study, circ-PGAP3 was shown to increase TNBC proliferation and invasion via miR-330-3p/Myc axis (45).

As a result of our study, many circRNAs, miRNAs and genes that may be associated with BC have been identified. We found that three circRNAs (hsa_circ_0016201, hsa_circ_0000375, hsa_circ_0000515), three miRNAs (hsa-miR-141-5p, hsa-mir-183-5p, hsa-miR-486-5p) and 18 genes (CIDEC, ADH1B, TMEM132C, ACVR1C, LIPE, ABCA8, BTNL9, TNXOCB3, GPAM, PRAME, MELK, NEK2, EXO1, TPX2, BUB1, DLGAP5, SDC1) may be important in BC, especially in basal-like group, by applying filters as described in the material method section. Afterwards, by using the bottleneck tool, SDC1 was detected as a hub gene.

Expressions of hsa-miR-141-5p and hsa-mir-183-5p, which are known to be dysregulated in many cancers including BC (46-50), were found to be significantly increased in our study in all dataset samples from all PAM50 groups. Possible target genes that may contribute to the cancer progression and in which hsa-miR-141-5p and hsa-mir-183-5p could alter their expression in BC are shown in the Table 2. According to the criteria we determined, the possible paired targets of hsa-miR-141-5p/ ADH1B and hsa-mir-183-5p/ BTNL9, may be related to the BC process. In the dataset we examined, it was identified that hsa_circ_0016201, which is among the circRNAs whose expression was significantly decreased, could have a role as a sponge for miR-141-5p and hsa_circ_0000375 could be acted as a sponge for miR-183-5p. Therefore, we would like to emphasize that the relationship between hsa_circ0016201/ miR-141-5p/ ADH1B and hsa_0000375/ miR-183-5p/ BTLN9 should be investigated at the cellular functional level. It could be substantial to examine this relationship with conventional molecular genetic techniques in both BC cells and tumor tissue.

More importantly the expression of hsa-miR-486-5p, which is an essential tumor suppressor miRNA in BC and many other cancer types (51-54), was significantly decreased in all PAM50 groups examined in our study. It was determined that hsa_circ_0000515, one of the circRNAs whose expression was significantly increased in the dataset we examined, could act as a sponge for miR-486-5p. In addition, we determined that the increased expression of SDC1, PRAME, EXO1, BUB1, and DLGAP5 genes could be more strongly associated targets of miR-486-5p in BC. The overexpressed SDC1 gene was found to lead a significantly poor OS and DFS and overexpressed PRAME, EXO1, BUB1 and DLGAP5 genes were found to lead a significantly poor DFS in BC (Fig. 9-11) (criterion 3-b). miR-486-5p has been reported as an important tumor suppressor miRNA in various cancers, including BC. It has been reported that miR-486-5p which could be found exosomal miRNA in BC inhibits epithelial-mesenchymal transition (EMT) by targeting Dock1 and suppresses cancer cell proliferation by targeting the PIM-1 oncogene in BC, can be used as a biomarker in the prediction of BC recurrence (53-55). There are valuable studies showing that miR-486-5p may be associated with different circRNAs. The importance of circNFIB1/miR-486-5p/PIK3R1/VEGF-C axis in pancreatic cancer lymphatic metastasis (56); hsa_circ_0016788/miR-486/CDK4 pathway in hepatocellular carcinoma tumorigenesis (57); circHUWE1/miR-486-5p in colorectal cancer migration and invasion (58) and Circ-TCF4.85/miR-486-5p/ ABCF2 in hepatocellular carcinoma progression (59) are reported in the literature. However, as far as we know the relationship between miR-486-5p and circRNA has not yet been reported in BC. Syndecan-1 (SDC1, CD138) is a critical cell surface adhesion molecule required for cell morphology and impact on the natural microenvironment. SDC1 dysregulation enhances cancer development by increasing cell proliferation, angiogenesis, invasion, and metastasis and is linked to chemo-resistance. SDC1 expression is also correlated to chemotherapy responses and prognosis in a variety of solid and hematological malignancies, including BC (60, 61). It has been suggested that SDC1 could be a new molecular marker that alters the phenotype of cancer stem cells through the IL-6/STAT3, Notch, and EGFR signaling pathways in triple-negative inflammatory BC (62). Induction of SDC1 in the lung microenvironment may promote the formation of breast tumor metastasis (63). SDC1 has been found to have a vital role in the progression of BC metastasis to the brain. SDC1 has been shown to increase BC cell migration across the blood-brain barrier via modulating cytokines, which may alter the blood-brain barrier (64). SDC1 overexpression in BC has been shown to be associated with various miRNAs (65, 66). However, the relationship between miR-486-5p/SDC1 and BC is not yet known. It has been reported that SDC1 expression can also be indirectly altered by circRNAs as it has been demonstrated that circCEP128 is associated with bladder cancer progression via the miR-515-5p/SDC1 axis (67). According to our bioinformatics study findings, we recommend further investigation of the SDC1 gene together with the hsa_circ_0000515/ miR-486-5p axis when conducting circRNA/miRNA/gene functional research in BC. We propose that hsa_circ_0000515/ miR-486-5p/ SDC1 axis may be an important biomarker candidate in distinguishing patients in the BLBC group, especially according to the PAM50 classification of BC.

## Conclusion

Finding new biomarkers to clearly classify subtypes of BC could be quite crucial in the battle against cancer. To identify novel biomarkers and new therapeutics, a deeper understanding of the mechanisms underlying BC metastasis is extremely important. According to our study results, we suggest various DE mRNAs, miRNAs and circRNAs that may be important in the onco-transcriptomic cascade for BC. The interrelationships of these molecules can be potential diagnostic biomarkers or therapeutic targets. Therefore, functional experiments such as proliferation, apoptosis, invasion, and metastasis on BC cells should be studied to elucidate these circRNA-miRNA-mRNA relationships in the future.

## Declaration of conflicting interest

There is no conflict of interest between the authors.

## Ethics approval and consent to participate

GEO, EGA and TCGA databases were used to download the data of this research.

## Funding

None

## Notes

### Competing Interest Statement

The authors have declared no competing interest.

